# Evolutionary origin of morphologically cryptic species imprints co-occurrence and sympatry patterns

**DOI:** 10.1101/2023.09.13.557531

**Authors:** Teo Delić, Špela Borko, Ester Premate, Behare Rexhepi, Roman Alther, Mara Knüsel, Florian Malard, Dieter Weber, Fabio Stoch, Jean-François Flot, Cene Fišer, Florian Altermatt

## Abstract

**Aim:** Morphologically cryptic species are an important part of global biodiversity, yet it remains unclear how these species contribute to and integrate into communities at different geographic scales. It is especially unclear at which scales they co-occur, and if and how their ranges overlap. To adequately protect biodiversity, an accurate understanding of the underlying processes and adequate level of protection is needed, in particularly for often overlooked cryptic species.

**Question:** We analyzed patterns of syntopies (local co-occurrences) and sympatries (range overlap) to test how the evolutionary origin of cryptic species shapes biodiversity patterns at different geographic scales. We hypothesized i) that syntopies are more common among phylogenetically deeply divergent cryptic species than among close relatives, and ii) that sympatries are an outcome of phylogenetic relatedness and dispersal, with range size as a proxy of dispersal ability.

**Location:** Western Europe.

**Taxon:** Subterranean amphipod crustaceans of the polyphyletic *Niphargus rhenorhodanensis* species complex.

**Methods:** Unilocus species delimitations (PTP, ASAP), calibrated multilocus phylogenetic analyses, co-occurrence analyses using a probabilistic model, generalized linear models (GLM).

**Results:** The studied species complex comprises 37–48 molecular operational taxonomic units (MOTUs) from nine different clades. Syntopies are random or less frequent than expected, implying an insufficient between-MOTU differentiation allowing stable coexistence. GLM suggested that age of divergence does not predict species sympatries, although they emerge more frequently among MOTUs from different clades. By contrast, sympatries emerge when at least one MOTU disperses over a large geographic range. Biodiversity rich regions were found at the foothills of the Alps, the Jura and the Central Massif, regardless the inference method.

**Main conclusions:** Biodiversity patterns of the herein studied species complex are driven mainly by dispersal and reflect geographic circumstances of speciation. While species richness on a local scale may be the outcome of competition and dispersal, regional biodiversity patterns emerged through biogeographic history on a clade-level.

## Introduction

Cryptic species are morphologically indistinguishable or near indistinguishable species that can be reliably delimited and recognized by molecular tools (Bickford et al., 2007). Cryptic species are increasingly found across all animal phyla and are an important part of local and global terrestrial and aquatic biodiversity (Eme et al., 2017; Li & Wiens, 2023; Pérez-Ponce de León & Poulin, 2016). In some taxa they represent only a minor fraction of species diversity, whereas in other they account for the majority of clade diversity (Adams, Raadik, Burridge, & Georges, 2014; Lukić, Delić, Pavlek, Deharveng, & Zagmajster, 2019; Wattier et al., 2020). While early authors assumed that morphological crypsis corresponds to ecological similarity, and that such species co-occur only exceptionally (Wellborn & Cothran, 2004), more recent studies indicate that co-occurrence of cryptic species can be common (C. Fišer, Robinson, & Malard, 2018; Henry et al., 2013; Marsteller, Adams, Collyer, & Condon, 2009; Voda, Dapporto, Dinc, & Vila, 2015). However, it is unclear how these species contribute to and integrate into communities across different geographic scales.

Community assembly and species’ co-occurrence are eventually shaped by speciation, dispersal, stochastic factors, and interspecific interactions (Vellend, 2010). The latter, interspecific interactions, deterministically affect if and how species can co-occur. Interspecific interactions are governed by a variety of factors, including ecological differentiation, and may also shape dispersal and evolution of different dispersal abilities. Modern coexistence theory suggests that the outcome of the between-species interaction is balanced by niche and ecological fitness differences (the latter approximated by population growth at low densities), resulting in species coexistence or exclusion, respectively (Chesson, 2000; Spaak & De Laender, 2020). On a local scale, ecologically different species coexist regardless the difference in their ecological fitness, whereas ecologically similar species may coexist only when both species have equal ecological fitness (unstable coexistence *sensu* (Chesson, 2000)). Such coexistences are only temporary and eventually end with exclusion of either of species, yet without predictable order. With increasing geographic scales, dispersal becomes more important in mediating community assembly mechanism by counterbalancing local extinctions and facilitating regional species coexistence through local colonization-extinction dynamics (Hart, Usinowicz, & Levine, 2017; Leibold & McPeek, 2006; Leibold, Urban, De Meester, Klausmeier, & Vanoverbeke, 2019; Luo, Wang, Saavedra, Ebert, & Altermatt, 2022). Here, we ask if the evolutionary origin of cryptic species constrains factors affecting species coexistence through either ecological differentiation, dispersal, or both, and hence affects patterns of species richness. We refer to coexistence on local and regional scales as “syntopy” and “sympatry”, respectively. Moreover, we use the phrase “patterns of species richness” when number of regionally coexisting species differ among regions.

Ecological differences contribute to coexistence on a local to regional scale. Ecological differences among cryptic species are hard to quantify, and do not correspond to morphological similarity (Eisenring, Altermatt, Westram, & Jokela, 2016). Arguably, ecological differentiation of cryptic species relates to their phylogenetic relatedness as phylogenetic contingency (Cavender-Bares, 2019; Cavender-Bares, Kozak, Fine, & Kembel, 2009; Dormann et al., 2018). If so, we firstly hypothesize that closely related (i.e., ecologically similar) cryptic species locally outcompete each other and at best establish sympatry, whereas deeply divergent (i.e., ecologically differentiated) cryptic species more frequently co-occur in syntopy and form broad areas of sympatry.

Dispersal can be triggered by interspecific interactions, such as predation and competition (Fronhofer et al., 2018; Little, Fronhofer, & Altermatt, 2019), but within the limits of a species range size also contributes to coexistence on a regional scale (realized dispersal (Malard et al., 2023). Species with large ranges inherently disperse over larger distances and thus contribute to community assembly on a regional level (Bregović, Fišer, & Zagmajster, 2019). Range sizes are an outcome of numerous factors (Borko, Premate, Zagmajster, & Fišer, 2023; Gaston, Trans, Lond, & Gaston, 1998), however, they initially depend on historical circumstances governing speciation. Speciation can fragment ancestral ranges into many narrowly distributed species (symmetric range splitting), or unevenly, where one or few species bud-off from the ancestral range, yielding at least one large-range species and one to many small-range species (asymmetric range splitting) (C. Fišer, Robinson, et al., 2018). Hence, symmetric *vs.* asymmetric range splitting at speciation may result in species with different realized dispersal, and thus indirectly affects community assembly on a regional scale. Thus, we secondly hypothesize that asymmetric range splitting during speciation enhances development of sympatries, because it results in species with larger ranges and higher degrees in realized dispersal.

Both hypotheses render two testable predictions. Firstly, we expect that closely related species pairs are ecologically more similar and thus form only unstable coexistences maintained through dispersal-outcompetition dynamics. The frequency of syntopic pairs predicted from occurrence frequencies of species-pairs will equal random expectation due to dispersal, or will fall below it due to competitive interactions. Conversely, phylogenetically divergent species pairs are ecologically differentiated, establish stable coexistence and, given enough time to disperse, co-occur in syntopy more frequently than expected from their frequencies of occurrences. Secondly, formation of sympatries depends mainly on dispersal whereas the importance of between-species interactions diminishes due to scaling effects. Within a region, species with large ranges contribute more to net dispersal than narrowly distributed species. Sympatries can be expected mainly among those pairs, where at least one species has a medium to large range, but not among narrowly distributed species that emerged through asymmetric range splitting during speciation.

Testing these predictions requires a species-rich complex of cryptic species, comprising species of different ages and overlapping ranges. The species complex of *Niphargus rhenorhodanensis* Schellenberg, 1937 fulfils these criteria. It belongs to a group of subterranean amphipods (Crustacea) widely distributed in Central to Southern Europe, counting at least ten species from minimally four lineages (Trontelj et al., 2009), whose ranges geographically broadly overlap (Eme et al., 2017; Lefébure, Douady, Malard, & Gibert, 2007). We amassed subterranean samples from hundreds of locations, analyzed the taxonomic structure of the *N. rhenorhodanensis* species complex using unilocus species delimitations, and examined the species’ distributions, range sizes as well as the phylogenetic structure within the species complex. This dataset allowed us to study how the mode of speciation and depth of phylogenetic divergence dictates the degree of sympatry and syntopy among cryptic species, and thus sheds light on how evolution of cryptic species contributes to community assembly over different geographic scales. We specifically i) tested whether pairwise species co-occurrences on a local scale deviate from random expectations; ii) analyzed whether distributional ranges of species pairs overlap with respect to their relatedness and range sizes; and iii) explored where sympatries peak and define species rich areas.

## Materials and methods

### The model system

Our model system is the species complex of *Niphargus rhenorhodanensis.* As a species complex, it covers a comparably large area for a groundwater organism (Trontelj et al., 2009), being widely distributed between the rivers Rhine and Rhône in Central to Southern Europe (occurring in parts of France, Switzerland, and Germany, but also extending to the catchment of the Po river in north-western Italy (Ginet, 1985; Karaman, 1993). Its morphological crypsis relates to high variation at the level of individual (ontogenetic variation), sex, within-and between population and prohibits a clear morphological diagnosis between species.

Based on molecular markers, however, *N. rhenorhodanensis* is not one species, but a complex of many morphologically similar, but genetically distinct species (C. Fišer, Alther, et al., 2018; Ginet, 1985; Lefébure et al., 2007; Trontelj et al., 2009). Morphology-based taxonomy was first challenged by allozymes (Berettoni, Mathieu, & Hervant, 1998), and then by molecular markers (Eme et al., 2017; Lefébure et al., 2007). Few morphologically distinct species of this species complex were recently described on a basis of morphology (C. Fišer, Konec, Alther, Švara, & Altermatt, 2017; Karaman, 2013), whereas a large majority of genetically distinct species remained undescribed because of their morphological similarity. It is clear that the complex is polyphyletic, and counts at least ten species (Trontelj et al., 2009). The different species’ ranges geographically broadly overlap (Eme et al., 2017; Lefébure et al., 2007). Hence, *N. rhenorhodanensis* is a species-rich complex of morphologically similar yet differently related species forming broad sympatries.

### Datasets and laboratory work

Over the past years, we collected samples of the *N. rhenorhodanensis* species complex from 151 localities from Switzerland, France, Germany, and Italy. These samples, along with already available sequences were the basis for three datasets, named *Delimitation*, *Phylogeny*, and *Spatial* datasets (Supplementary Material 1–3).

The *Delimitation* dataset was used for inferring the number of putative species within *Niphargus,* including the herein studied species complex. We gathered all *Niphargus* COI sequences from GenBank and supplemented it with our own unpublished data available until April 2022. After checking and cleaning the dataset, we ended up with 3,310 individuals that were subjected to species delimitations. Of these, 764 individuals were sequenced *de novo*. Detailed information is available in Supplementary Material 1. The results derived from this dataset, molecular operational taxonomic units (MOTUs), were used as proxies for species in the subsequent analyses.

The *Phylogeny* dataset was used to infer the phylogenetic structure of the *N. rhenorhodanensis* species complex within the genus *Niphargus*, and to assess how many clades it comprises. We assembled a dataset of 561 MOTUs. These were sequenced for up to seven markers, including mitochondrial COI, two segments of the 28S rDNA, histone gene (H3), partial sequence of the 18S rRNA, partial sequence of the glutamyl-prolyl-tRNA synthetase gene (EPRS), heat shock protein 70 gene (HSP70), and arginine kinase (ArgKin) gene, as specified in (Borko, Altermatt, Zagmajster, & Fišer, 2022; Borko, Trontelj, Seehausen, Moškrič, & Fišer, 2021). For the needs of the study, 382 sequences were obtained *de novo*. Detailed information is available in Supplementary Material 2.

The *Spatial* dataset was used for spatial analyses where we evaluated the frequency of syntopies (local co-occurrences), degree of sympatry (range overlap, regional scale) and geographic pattern of MOTU richness. The dataset comprised all individuals identified as *N. rhenorhodanensis*, and already described species nested within this complex (*Niphargus catalogus, Niphargus styx*). This resulted in 376 individuals of the herein studied species complex, collected at 144 localities. Detailed information is available in Supplementary Material 3. In total, we extracted and amplified DNA from 764 individuals in three laboratories (SubBio Lab, Ljubljana, Slovenia; Eawag, Dübendorf, Switzerland; EEB, Brussel, Belgium). The specifics of the laboratory protocols are separately given in Supplementary Material 4.

### Species delimitation and estimation of number of species

We employed two broadly used species delimitation methods, ASAP (Assemble Species by Automatic Partitioning) (Puillandre, Brouillet, & Achaz, 2021) and PTP (Poisson Tree Processes) (Zhang, Kapli, Pavlidis, & Stamatakis, 2013). Although comparative analysis of delimitation methods unveiled that ASAP returns the most reliable results on large datasets (Borko et al., 2022), we ran PTP as a control.

The ASAP delimitation was executed under the Kimura two-parametric test (Kimura-K80), with a prior for maximum value of intraspecific divergence between 0.005 and 0.5, ten recursive steps and a gap width of 1.0. We executed ASAP delimitation using the complete dataset on an online server available at https://bioinfo.mnhn.fr/abi/public/asap/.

For the tree-based PTP, we first removed duplicate haplotypes using a custom Perl script (Eme, Malard, Konecny-Dupré, Lefébure, & Douady, 2013). The remaining haplotypes (N = 1,982) were then used to infer phylogenetic relationships using IQtree 1.6.7 (Nguyen, Schmidt, Von Haeseler, & Minh, 2015) by deploying codon specific substitution models. The resulting trees were then used to run the PTP analysis on the species delimitation server https://mptp.h-its.org/#/tree under the default settings.

The two analyses yielded two alternative groups of MOTUs. These were subjected to comparative examination with published data from multilocus and integrative taxonomic studies, as described previously (Borko et al., 2022), and the finally accepted MOTUs were treated as species proxies in subsequent phylogenetic and spatial analyses. Details on the samples are available in Supplementary Material 1.

To evaluate how well sampling effort covered the species structure of the *N. rhenorhodanensis* species complex, we statistically estimated species richness from the available samples. Using MOTUs obtained with a more conservative ASAP, we analyzed how the number of MOTUs increases with addition of samples. To do this, we ran a rarefaction curve using 1,000 random runs and estimated the asymptote. Additionally, we calculated Chao2, an index that estimates species richness based on the number of all species corrected for rare species, i.e., species found only once (uniques) or twice (duplicates) in the samples. For the analyses, we used EstimateS (Version 9.1.0) (Colwell, 2013) based on the *Spatial* dataset (Supplementary material 3).

### Phylogenetic analyses

Phylogenetic analyses were based on the above mentioned species delimitations, and followed the procedure used in two previous studies (Borko et al., 2022, 2021). Briefly, we calculated a multilocus time-calibrated tree using 560 available MOTUs of *Niphargus* and one outgroup (Supplementary Material 2). For each marker, we aligned the sequences in Geneious 11.0.3 (Biomatters Ltd, New Zealand), using MAFFT v7.388 plugin (Katoh & Standley, 2013), with the E-INS-I algorithm with scoring matrix 1PAM/k=2 and the highest gap penalty. We concatenated alignments in Geneious. We removed gap-rich regions from the alignments of non-coding markers using Gblocks 0.91.1 (Talavera & Castresana, 2007) with a less stringent selection set. The optimal substitution model for each partition was inferred using Partition Finder 2 (Guindon et al., 2010; Lanfear et al., 2012) under the corrected Akaike information criterion (AICc).

A time-calibrated chronogram was built using BEAST 2 2.7 (Bouckaert et al., 2014), using four congruent calibration points referring to *Niphargus* fossils from Eocene Baltic amber (corresponding to the first occurrence of pectinate dactylus, 40.0–50.0 Mya), the age of the final submergence of the land bridges between Eurasia and North America (root age, 71.0–57.0 Mya), the age of the last land-bridge between Crete and Greek mainland (age of Crete species, 5.3–5.0 Mya) and the opening of the connection between Paratethys and the Mediterranean basin (age of the Middle East clade derived from Eastern European taxa, 11.0–7.0 Mya). Sensitivity analysis to test congruence among calibration points and additional details are available in (Borko et al., 2021). For each gene partition, we employed the following set of priors: linked birth-death tree model, unlinked site models with fixed mean substitution rate, and relaxed clock log-normal distribution with estimated clock rate. We used default settings of distributions of all estimates. We ran the analyses for 200 million generations, sampled every 40,000 generations. The first 25 % of the trees were discarded as burn-in and results were inspected in Tracer 1.7 (Rambaut, Drummond, Xie, Baele, & Suchard, 2018).

### Syntopies, sympatries and biodiversity patterns

In spatial analyses, we used MOTUs delimited by ASAP, which returned more conservative species delimitations hypothesizing fewer species. Range sizes of MOTUs were estimated using maximum linear extent (MLE), a linear distance between the two most distant localities of a MOTU (using projection EPSG: 5627). This measure was commonly used in analyses of range sizes of groundwater organisms (Borko et al., 2023; Bregović et al., 2019; Zagmajster et al., 2014).

To test MOTUs co-occurrences within each locality (syntopies), we employed a probabilistic model of species co-occurrence with hypergeometric distribution. The model tests the observed pairwise MOTU co-occurrences against the expected MOTU occurrences, calculated from the empirical data (Veech, 2014). The model returns whether MOTU pairs co-occur at random (as expected with respect to species rarity), more frequently than expected, or less frequently than expected. In these analyses we included also two described species (*N. styx, N. catalogus*) as they are nested within the species complex (see Results).

For the analyses, we used the *Spatial* dataset. To minimize violation of the assumption of unlimited dispersal, we split the dataset into two, geographically distinct sub-datasets separated by the Alpine arch, comprising localities from Italy (hereafter “Mediterranean”), France, Switzerland and Germany (the latter three hereafter “Continental”). We removed one very remote locality in the northwest of France from the “Continental” subset, as it is highly dispersal limited and isolated. For each sub-dataset, we asked whether MOTUs pairwise co-occurrences deviate from random. Although we sequenced multiple individuals from each locality, we cannot rule out false negative records of co-occurrences when one species is rare. To account for the potential bias, we repeated the analyses using potential co-occurrences in a radius of 5 km.

To analyze between-MOTU sympatries, we first defined the geographic distributions of individual MOTU as minimum convex polygons. MOTUs known from a single locality were assigned a distribution with a diameter of 10 km. We defined a pair of MOTUs sympatric when their ranges (minimum convex polygons) overlapped. We filtered sympatric MOTU pairs from all possible pairs and first examined whether sympatric pairs comprise species from the same clade (ecologically more similar), or from different clades (ecologically differentiated). To test whether sympatries emerge as frequently among members of the same clades as among members of different clades, we employed a χ^2^-test. Additionally, we tested whether sympatry is more frequent between phylogenetically more distant MOTUs using a generalized linear model with a binomial distribution. All MOTU pairs were assigned a dummy variable, whether or not they form a sympatry, and divergence times, i.e., time since the last common ancestor. From all possible MOTU pairs, we removed unrealistic pairs of species living on both sides of the Alpine arch (see above). To model how phylogenetic distance predicts pairwise sympatries, we ran a generalized linear model using binary defined sympatry-nonsympatry as response variable, and divergence times as predictors.

Finally, to test whether range size predicts pairwise sympatries, we ran another generalized linear model with a binomial distribution, using range sizes as predictor and binary variable (sympatry-nonsympatry) as response variable. Given that MOTUs have different range sizes, we summed MLEs for each MOTU pair. Again, we excluded trans-Alpine pairs that likely violate the assumption of unrestricted dispersal. The data used in both generalized linear models are available in Supplementary Material 5. We also evaluated at what altitudes peak MOTU and clade diversity. To do this, we quantified the surface of polygons of overlapping MOTU / clade ranges. Each polygon was overlaid by a grid of 25 x 25 m resolution using SAGA: resample (in QGIS) and mean value (cell area weighted) as upscaling method. Input rasters: EU-DEM v1.1 maps E30N20 and E40N20, Copernicus Land Monitoring Service https://land.copernicus.eu/imagery-in-situ/eu-dem/eu-dem-v1.1. Extraction of cell values within areas made in R using raster (Hijmans 2023).

We ran the analyses in R 4.2.2 (R Development Core Team, 2022) and RStudio 2023.06.0 (RstudioTeam, 2023) using packages *cooccur 1.3* (Griffith, Veech, & Marsh, 2016) and *sf 1.0-9* (Pebesma, 2018), and IBM SPSS version 20. We produced the figures using R packages *ggplot2 3.4.0* (Wickham, 2016) and *alluvial 0.1-2* (Bojanowski & Edwards, 2016). We created all maps in QGIS 3.10.12 (QGIS.org, 2020) and phylogenetic trees in FigTree 1.4.3. (Rambaut, 2016).

## Results

### Species delimitation

Overall, species delimitations applied to the entire genus *Niphargus* yielded 491 (ASAP) and 679 (PTP) MOTUs. Of these, 37 ASAP and 48 PTP delimited MOTUs belonged to the studied complex *N. rhenorhodanensis* s. lat. (Table S2 in Supplementary Material 7). Two species within the herein studied complex have been described as *N. styx* and *N. catalogus* (Supplementary Materials 1 and 3) (C. Fišer et al., 2017; Karaman, 1995). The difference in taxonomic structure between the two methods derives from the cut-off level at which either method recognizes species boundaries and not from a mis-matched grouping of individuals.

The species rarefaction curve based on MOTUs delimited by ASAP indicated that sampling efforts should be approximately three times higher to reach the asymptotic 57 species (38–77, 95 % confidence interval, Supplementary Material 8, Fig. S2). Asymptotically determined species richness was in the same range as species richness estimated by Chao2 (66, species, 45–130, 95 % confidence interval). In any case, the species richness of the studied cryptic species complex may be about two times higher than empirically observed.

### Phylogenetic relationships

Phylogenetic analysis recovered *Niphargus* relationships described in previous studies (Fig. S1 in Supplementary Material 5 compared to (Borko et al., 2022, 2021)), with somewhat increased node support. The phylogenetic structure suggested that *N. rhenorhodanensis* comprises nine clades (Fig. 1), separated at least 10 Mya, which we consider independent from each other (hereafter clade 1–9). Several clades are nested in different parts of *Niphargus* phylogeny and imply that some morphological similarity evolved by convergence. Each clade counts between one and 12 MOTUs. Five clades comprise more than one MOTU. The taxonomic structure, age, MLE and geographic distribution of the five major clades are summarized in Table 1, Fig. 2 and Fig. S3 (Supplementary information 8). Analysis of cladogenetic events within five clades suggested mostly asymmetric speciation events, where each clade comprises one or few large-ranged species and several narrowly distributed species. Species with narrow distributional ranges proliferated mainly in montane regions (Fig. 4).

**Fig. 1.**
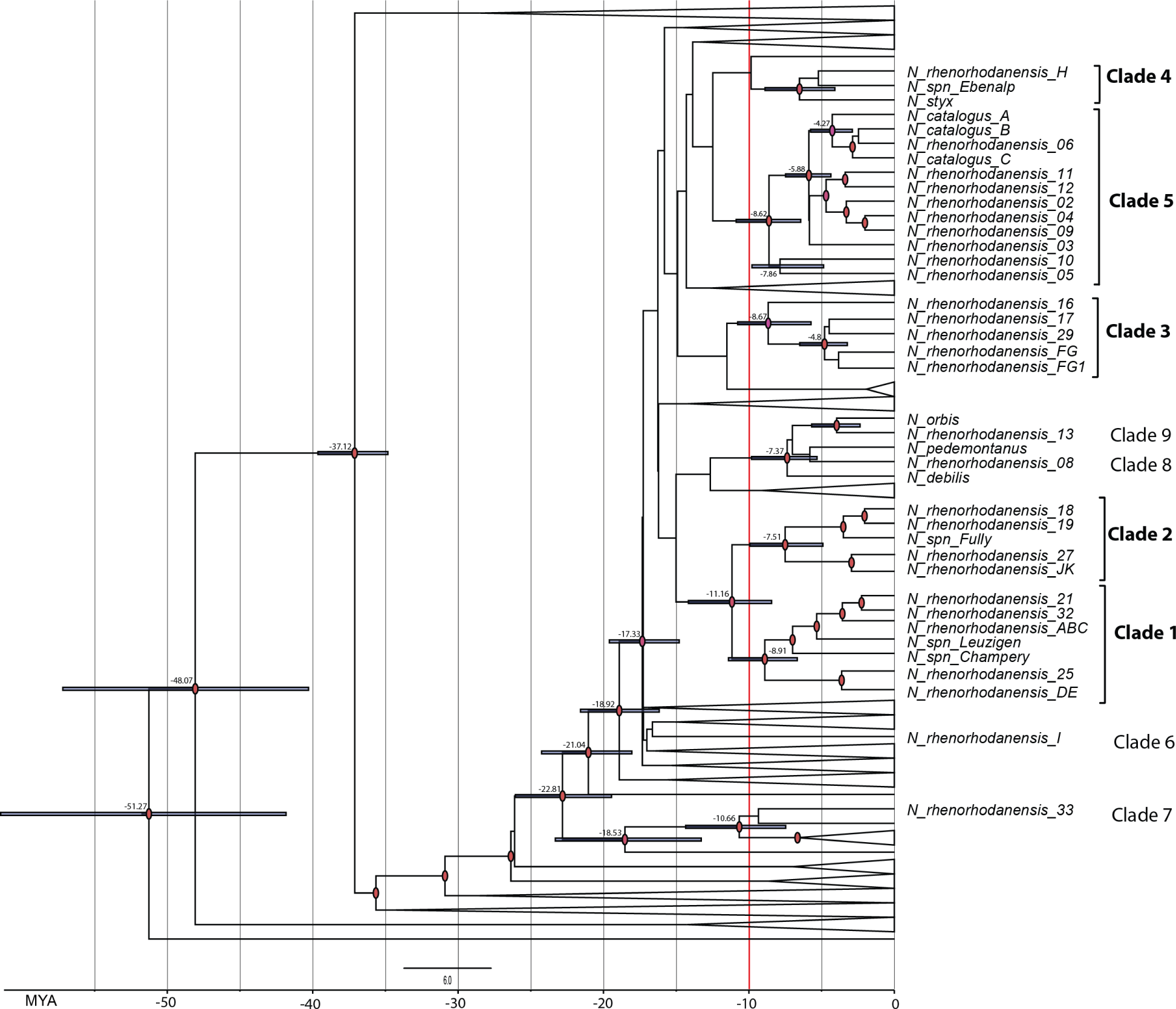
Calibrated phylogeny of 561 MOTUs obtained by Bayesian inference and phylogenetic structure of the *N. rhenorhodanensis* species complex. We simplified the phylogeny by removing outer clades, collapsing inner clades, and retaining the age only on some of the nodes. The threshold of 10 MY is highlighted by a red vertical line. Posterior probabilities above 0.9 are indicated with red ellipses at the node. A complete phylogeny is available in Supplementary Material 6, Fig. S1.

**Table 1.** A summary of combined results from species delimitation and phylogenetic analysis. Old labels refer to MOTU labels used in previous publications (Borko et al., 2022; Lefébure et al., 2007). Age presents 95 % credibility interval. “Described species” indicates species that are part of a clade and were taxonomically described and named before this study.

| Clade | Old labels | No. MOTU | No. individuals | Age (MY) | Clade distribution | Described species |
| --- | --- | --- | --- | --- | --- | --- |
| 1 | ABCDE | 7 | 118 | 8.91<br>(5.36–14.56) | Central Massif, south of Jura and W Germany | na |
| 2 | JK | 5 | 16 | 7.51<br>(3.72–15.23) | Switzerland | na |
| 3 | FG | 5 | 128 | 8.67<br>(5.1–17.66) | Jura, Rhone valley and Central Massif | na |
| 4 | H | 3 | 89 | 6.52<br>(2.9–11.37) | Central Massif, Rhone Valley, Jura, Rhine Valley, inner Switzerland | *N. styx* |
| 5 | NA | 12 | 20 | 8.62<br>(4.79–13.98) | Italy | *N. catalogus* |

**Fig. 2.**
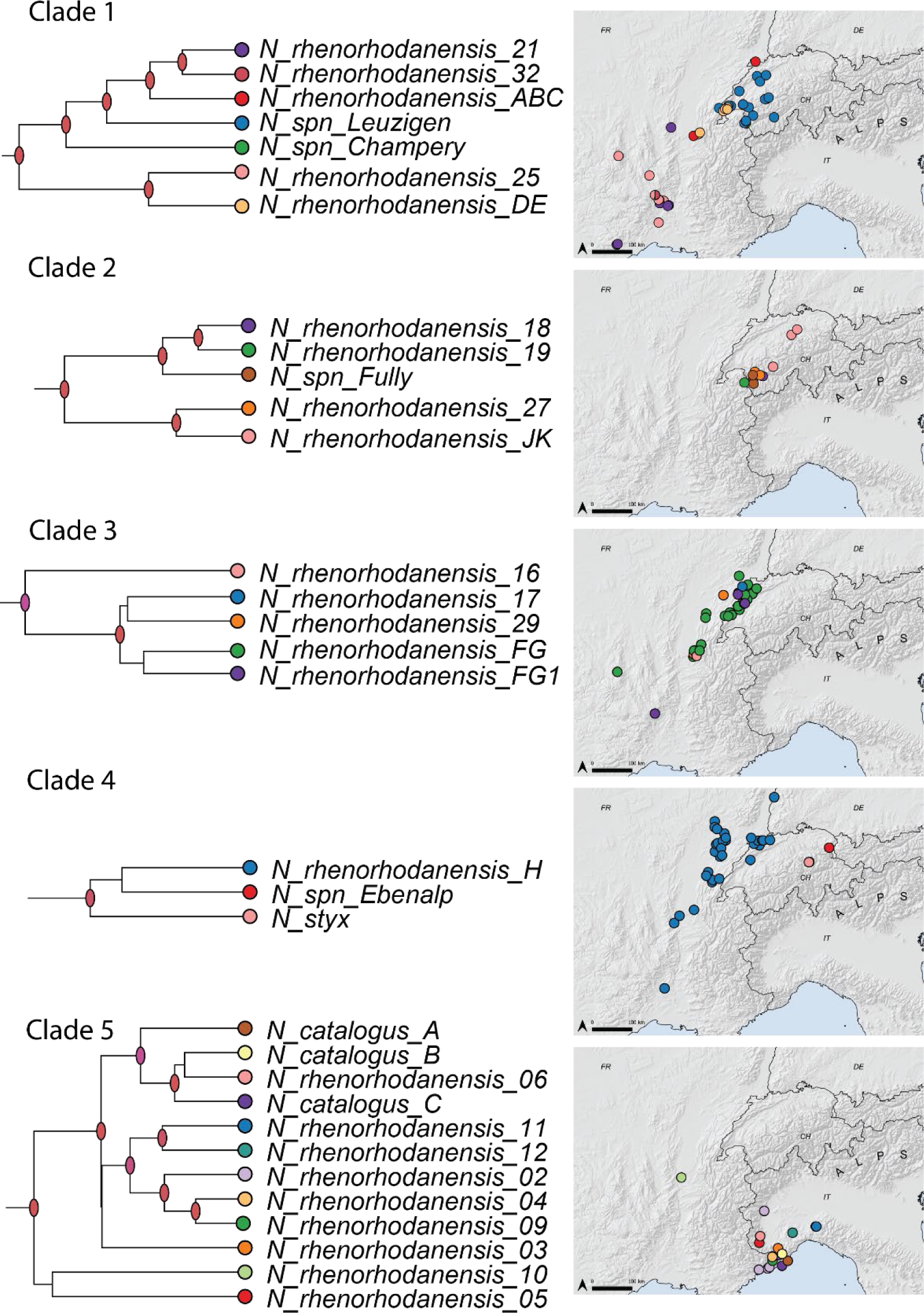
Five main clades and MOTU distribution. Most of the MOTU diversity of the studied complex is aggregated in five major clades. Individual clades are extracted from Fig. S1. Note that each clade contains few broadly and few narrowly distributed species.

### Syntopies, sympatries and biodiversity patterns

The spatial extent of individual MOTUs varies from a single site to several hundred kilometers along its longest axis. Clades 1, 2, 3, 4, and 5 comprise five, one, three, one, and one MOTUs with MLE > 100 km (Fig.2, Supplementary Material 5, Supplementary Material 8, Fig. S3).

Analyses of syntopies implied that co-occurrence records did not deviate from random expectation, or that co-occurrences were less frequent than expected from random, regardless the region we studied (Table 2). When we considered a potential bias due to false negative records, the result did not change in the Mediterranean region, whereas in the Continental dataset approximately 20 % of pairs co-occurred less frequently than expected. Among these, 11 pairs comprised members of the same clade and 39 pairs comprised members of different clades (Table 2).

**Table 2.** Results of co-occurrence analysis. We separately analyzed co-occurrences in France-Switzerland-Germany (Continental subset) and co-occurrences in Italy (Mediterranean subset), which are separated by the Alpine arch and violate an assumption of unlimited dispersal. The analyses were repeated on a level of MOTU and clade, i.e., where MOTU identity was replaced by clade membership. Each analysis was repeated at two geographic resolutions, one on exact locality and one using buffer zone of 5 km, to account for false negative records.

| geographic precision | region | Number of co-occurring pairs | Pairs co-occurring significantly less frequent than expected | Pairs co-occurring at random | Pairs co-occurring significantly more frequently than expected |
| --- | --- | --- | --- | --- | --- |
| exact locality | Continental (France, Switzerland, Germany) | 253 | 3 | 249 | 1 |
|  | Mediterranean (Italy) | 66 | 0 | 66 | 0 |
| Buffer zone with 5 km radius | Continental (France, Switzerland, Germany) | 253 | 50 | 221 | 4 |
|  | (Mediterranean (Italy) | 66 | 0 | 64 | 2 |

After removal of geographically remote MOTUs and separation of trans-Alpine pairs, we identified 356 possible MOTU pairs, of which 36 overlap in their ranges. Sympatries emerged among large-range MOTU pairs (MLE > 100 km, 20 pairs), or among large-small range MOTU pairs (16 pairs), resulting in statistically significant generalized linear model (test of model effect: Wald χ^2^ = 50.6, df = 1, p < 0.001, Fig. S4) (Fig. 2). These sympatries seem to be more common among MOTUs from different clades than among MOTUs of the same clade, yet with only marginally significant statistical support (Fig. 3, χ^2^ test, p = 0.077). Sympatric pairs diverged between 1.2–17.2 Mya. The results of the generalized linear model suggest that time since between MOTUs divergence does not predict origin of sympatry (phylogenetic distance GLM, p = 0.66, Fig. S5).

**Fig. 3.**
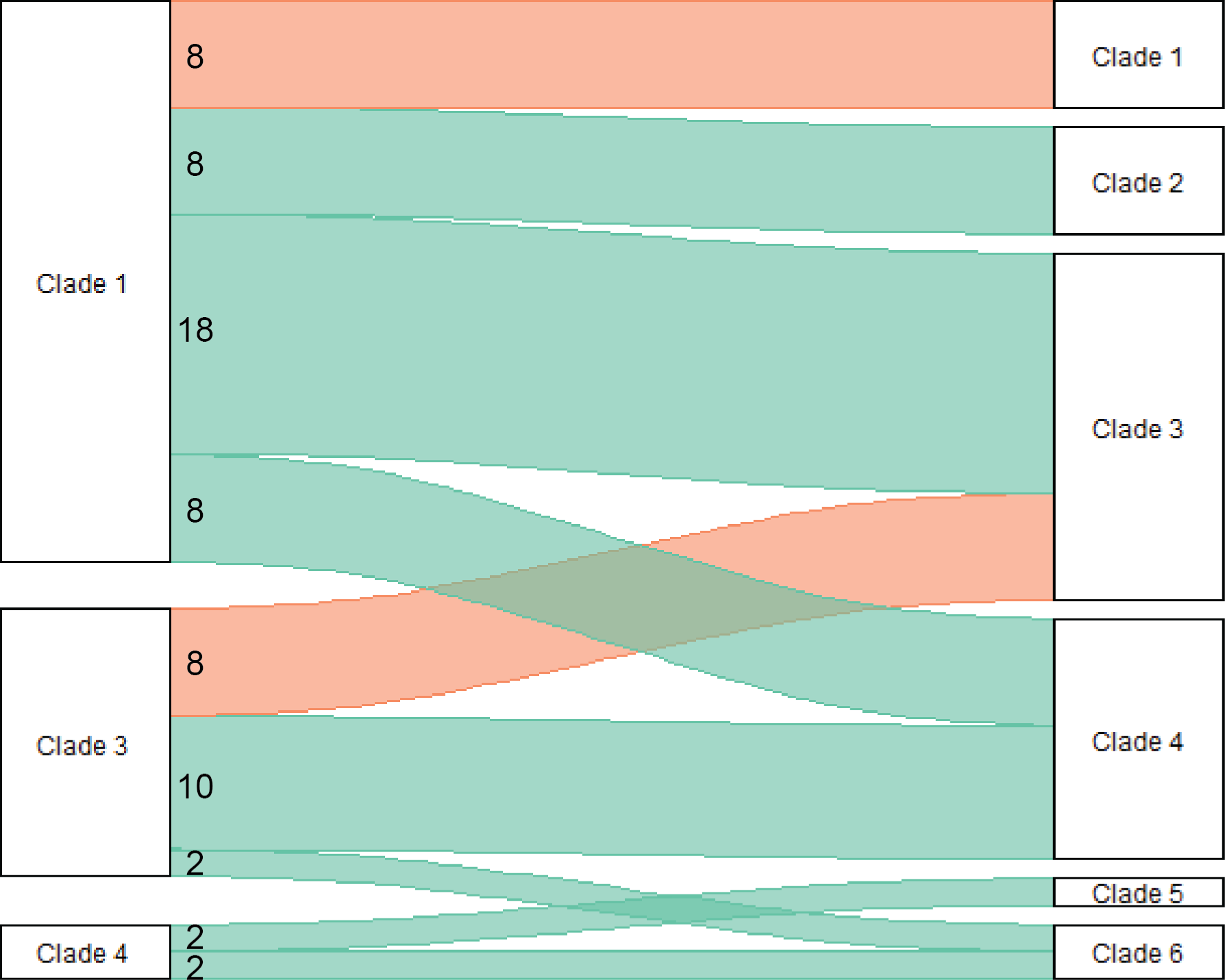
Schematic presentation of sympatric MOTU pairs belonging to the same or different clade. Overall, we identified 36 sympatric MOTU pairs from the *N. rhenorhodanensis* species complex. Most of these pairs derive from different clades (blue colored connective bands) and less so from the same clade (red colored connective bands). The width of a band correlates with the number of sympatries on a right side.

Overall, MOTU richness of the *N. rhenorhodanensis* complex peaks at the junction of the Jura Mountains and the Alpine arch, close to the political boundary between France and Switzerland, where up to six MOTUs overlap in their distributional ranges. Another area of higher MOTU richness falls into the Central Massif (France), where up to four MOTUs overlap in their ranges. We also checked overlapping ranges of individual clades (i.e., clade members treated as a single taxon), and regions of high clade diversity roughly correspond to regions of high MOTU diversity (Fig. 4). Regions, in which overlap most of the MOTU or clade ranges are at relatively low altitudes, between 200 and 500 m above sea level (Fig. 4, Fig S6 and S7).

**Fig. 4.**
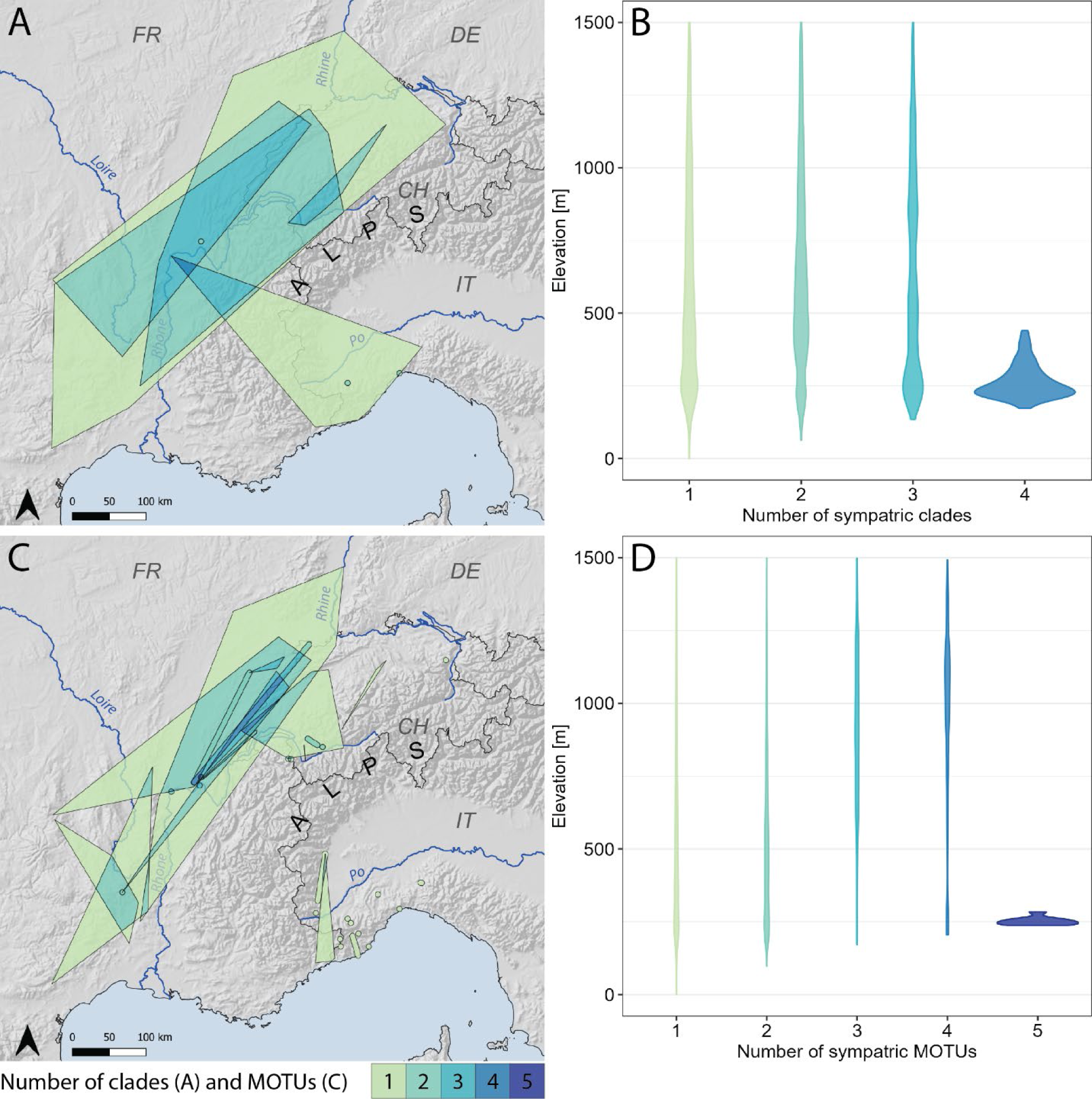
Biodiversity pattern formed by species of the *N. rhenorhodanensis* species complex. We overlaid minimal convex polygons of clades 1–5 (A) or MOTUs (C), respectively. Darkness of the layers corresponds to clade or MOTU richness, respectively. Note that clade- and MOTU-level imply similar regions of species richness, which peaks along the mountain foothills in the Central Massif, the Jura, and the Alps. Violin plots (B, D) represent elevation frequencies of 1 x 1 km squares within the areas covered by sympatric clades (B) and MOTUs. Note that we truncated plots at the elevation of 1500 m a.s.l.; plots showing the entire elevational range are available in supplementary material as Figs. S6–7.

## Discussion

We found clear signals of evolutionary origin on contemporary ecological patterns and community assembly of groundwater amphipods of the *Niphargus rhenorhodanensis* complex at different geographic scales, although not always matching our initial predictions. On a local scale, MOTUs of the studied cryptic complex were found to either co-occur at random, or less frequent from random expectation. This pattern is robust regardless of the species’ phylogenetic divergence and was detected in previous studies (Delić, Trontelj, Rendoš, & Fišer, 2017; Ž. Fišer, Altermatt, Zakšek, Knapič, & Fišer, 2015). This pattern may have emerged as a result of metacommunity dynamics, where members of different MOTUs come into contact with each other, temporarily coexist but eventually one species excludes the other one, either competitively or purely by chance. This could indicate these species are ecologically similar and functionally redundant, as competitive exclusion is an outcome of insufficient differences in ecological niches and ecological fitness (Luo et al., 2022). However, this pattern can also be partly explained with an alternative model, namely that early colonizers rapidly adapt to local conditions and impede subsequent colonization of preadapted newcomers through priority effects (Leibold et al., 2019). The latter hypothesis possibly better applies to pairs of species having large and small rages. Realized dispersal of species with small ranges seems to be limited, hence they need to have higher ecological fitness to exclude competing newcomers from neighboring regions (i.e., large-range species). While our comparative macroecological approach does not allow to establish causalities, we cautiously conclude that proliferation of cryptic species minimally affects species richness at the local scale (Delić, Trontelj, et al., 2017). This conclusion seems robust regardless the evolutionary mechanism behind the origin of cryptic species.

Sympatric MOTU pairs always comprise at least one MOTU with MLE exceeding 100 km, indicating that dispersal becomes important in community assembly processes at regional scale. Possibly, historical circumstances of speciation yielding species with different distributional ranges, contribute to sympatry formation. An illustrative case is Clade 2. Although the range of this clade overlaps with ranges of Clades 1, 3 and 4 (Fig. 2 and 4), its narrowly distributed members rarely form sympatries with other species (Fig. 3). Sympatry formation, however, is a more complex process also including possible roles of ecological differentiation inferred from the between-MOTU phylogenetic divergences. Although time since speciation alone does not explain sympatry origin (see Fig. S5), there is a weak tendency that phylogenetically unrelated cryptic MOTUs from different clades form sympatries (and syntopies) more frequently than closely related ones, possibly facilitating range overlap (Fig. 3). If true, this tendency is in line with our initial premise that phylogenetically unrelated species differ in ecological and physiological traits independently to morphological similarity (e.g. (Delić, Švara, Coleman, Trontelj, & Fišer, 2017; Eisenring et al., 2016; C. Fišer & Koselj, 2022; Scriven, Whitehorn, Goulson, & Tinsley, 2016). Such differences could facilitate local adaptation in a network of slightly different habitat patches, and mediate regional coexistence through spatial storage (Chesson, 2000; Hart et al., 2017) or priority effects (Leibold et al., 2019). Hence, sympatries among cryptic species likely emerge as a combination of how the range splits during speciation (asymmetric *vs.* symmetric) and the phylogenetic origin of cryptic species (members of the same clade versus different clades).

Finally, we observed characteristic patterns of highest MOTU richness at mountain foothills. This pattern can have at least two mutually non-exclusive explanations. Firstly, species along mountain ranges avoid extinctions by vertical dispersal and maintaining their ecological niches even during climatically unstable periods. Alternatively, it could indicate ecological gradients, possibly enforced by Pleistocene glaciations, prompted speciation through so called “speciation pumps”(Hu et al., 2021), and species rich regions emerge at the junction of species from different elevations. Intensified speciation at mountain slopes is well-known for freshwater amphipods in general (Copilaş-Ciocianu & Petrusek, 2015; Liu et al., 2023; Mamos, Wattier, Majda, Sket, & Grabowski, 2014) and *Niphargus* in particular (Delić et al., 2021; Lefébure et al., 2007; Stoch, Salussolia, & Flot, 2022). Interestingly, the map showing the biodiversity pattern of MOTUs is similar to the map showing biodiversity pattern of clades (Fig. 4). This corresponds to the above observation that patterns of MOTU diversity are driven by large-range MOTUs, which comprise most of the clade distributional range. It also suggests that the biodiversity pattern changed little since the onset of individual clade’s emergence 10–7 Mya, and that was possibly driven by climate changes during mid-late Miocene. Late Miocene was a period of climate cooling, reorientation of dominant winds and overall aridification (Kürschner, Kvaček, & Dilcher, 2008; Miao, Herrmann, Wu, Yan, & Yang, 2012) but see (Quan, Liu, Tang, & Utescher, 2014). The disputed aridification could have influenced hydrographic regimes, groundwater level and connectivity between ancestral habitats, yielding an extensive fragmentation of ancestral ranges and origin of the different clades. Whatever prompted speciation at that time, more recent speciation events only little contributed to overall biodiversity patterns. These findings are consistent to findings of Eme and coauthors (2017), who noted that biodiversity patterns on a large scale are relatively insensitive to the presence of cryptic species. Overall, we propose that historical biogeographic events, that is, the origin from different clades and asymmetric range splitting at the time of speciation, greatly shaped regional patterns of pairwise sympatries and MOTU-rich regions, whereas low MOTU diversity on a local scale is explained better by low ecological differentiation and local competitive exclusion.

Our study requires some cautionary notes. We used unilocus-delimited MOTUs as a proxy of species, given that the genus *Niphargus* has still many formally undescribed species. The use of MOTUs can be prone to oversplitting (Hupało, Copilaș-Ciocianu, Leese, & Weiss, 2022). To reduce such an effect, we built our analyses using conservative delimitation approaches. Specifically, the ASAP delimitation lumped several MOTUs delimited by PTP, and even some MOTUs delimited by ASAP using a smaller dataset (parallel analysis, data and results not shown). Previous comparison with multilocus delimitations and integrative taxonomy indicates that ASAP yields fairly accurate delimitations in *Niphargus*, when using large datasets (Borko et al., 2022). We are thus confident our delimitation analysis minimized the risk of oversplitting within the *N. rhenorhodanensis* species complex. Additionally, while extensive, our dataset may not yet completely cover the contemporary taxonomic diversity, and miss out some species that would be only found with increasing sampling efforts. Such additional species/occurrences would yield new co-occurrence records, and possibly modify species richness patterns. Finally, while extensive, our sampling was not systematically covering all areas with equal efforts. As inherent to such datasets, some regions had more intense sampling. Some MOTU were found only on some localities and have narrow distributional ranges. In case these species lived in subterranean compartments that we were not able to sample, their actual range size is underestimated. This could underestimate number of syntopies and sympatries. Yet, given that we sampled species using different equipment and different sampling approaches, and use here a compilation of data acquired by multiple independent research groups and sampling campaigns, we minimalized the risk of sampling bias affecting our results.

While we focus on the specific model system of groundwater amphipods, it allows to address questions of general interest on how speciation and adaptive evolution affect biodiversity patterns. *Niphargus* evolved in a series of adaptive radiations (Borko et al., 2021) originating several distinct ecomorphs (Delić, Trontelj, Zakšek, & Fišer, 2016; Trontelj, Blejec, & Fišer, 2012), as is also known from other adaptive radiations. The herein studied *N. rhenorhodanensis* species complex belongs to a slender, short-legged and sexually dimorphic ecomorph. Cryptic species were detected in many other ecomorphs (Delić, Švara, et al., 2017; McInerney et al., 2014; Švara, Delić, Rađa, & Fišer, 2015; Trontelj et al., 2009; Zagmajster & Fišer, 2009) as well. Different ecomorphs live in groundwater habitats with different connectivity and may have thus different range sizes (Borko et al., 2023). Hence, it may be expected that cryptic diversity from different ecomorph classes yield different biodiversity patterns, which eventually overlay one over another. A possible candidate for such an overlaid and complementary cryptic diversification of a functionally different clade is the cryptic *Niphargus virei* species complex with a different body shape and ecological properties (Foulquier, Malard, Lefebure, Douady, & Gibert, 2008; Issartel et al., 2006; Lefébure et al., 2006). It would be interesting to compare the phylogenetically distinct yet spatially overlaying biodiversity patterns of the *N. virei* and *N. rhenorhodanensis* species complexes, to address if and how patterns of coexistence and diversity within a complex are mirrored in other complexes too (pointing to large-scale drivers) or if they are different (pointing to more stochastic dynamics). Eventually, this can be framed as a more general question, namely how different evolutionary processes, leading to different ecomorph classes and complexes of cryptic species, jointly affect biodiversity patterns. Both, adaptive radiations (Gillespie et al., 2020; Schluter, 2000) and cryptic diversity (C. Fišer, Robinson, et al., 2018; Li & Wiens, 2023) are common phenomena and call to be integrated into macroecology (Eme et al., 2017).

## Acknowledgments

This research was funded within the DarCo project by Biodiversa+, the European Biodiversity Partnership under the 2021-2022 BiodivProtect joint call for research proposals, co-funded by the European Commission (GA N°101052342) and with the funding organisations Ministry of Universities and Research (Italy), Agencia Estatal de Investigación – Fundación Biodiversidad (Spain), Fundo Regional para a Ciência e Tecnologia (Portugal), Suomen Akatemia – Ministry of the Environment (Finland), Belgian Science Policy Office (Belgium), Agence Nationale de la Recherche (France), Deutsche Forschungsgemeinschaft e.V. – BMBF-VDI/VDE INNOVATION + TECHNIK GMBH (Germany), Schweizerischer Nationalfonds zur Förderung der Wissenschaftlichen Forschung (Switzerland), Fonds zur Förderung der Wissenschaftlichen Forschung (Austria), Ministry of Higher Education, Science and Innovation (Slovenia), and the Executive Agency for Higher Education, Research, Development and Innovation Funding (Romania). TD, BR, ŠB, EP and CF were additionally supported by Slovenian Research Agency (core program P1-0184, project J1-2464, PhD grants to Ester Premate and Špela Borko). FA, MK and RA were supported by grants by the Swiss Federal Office for the Environment (BAGU/FOEN, grant “Amphipod.CH”), the Swiss National Science Foundation (grants 310030_197410 and 31BD30_209583) and the University of Zurich Research Priority Programme on Global Change and Biodiversity (URPP GCB) to Florian Altermatt. F.M. was supported by the French National Research Agency projects CONVERGENOMICS (ANR-15-CE32-0005) and EUR H20’Lyon project (ANR-17-EURE-0018). Part of the equipment used in this study was purchased for the project “Development of research infrastructure for the international competitiveness of the Slovenian RRI space –RI-SI-LifeWatch”. The operation is co-financed by the Republic of Slovenia, Ministry of Education, Science and Sport and the European Union from the European Regional Development Fund.

