## SupplementaryMaterial04 for "Evolutionary origin of morphologically cryptic species imprints co-occurrence and sympatry patterns"

**Supplementary information 4**

**Laboratory Work**

For the needs of this study, we extracted and amplified DNA from 752 individuals using either GenEluteMammalian Genomic DNA Miniprep Kit (Sigma-Aldrich, United States) or MagMAX DNA Multi-Sample Kit (Thermo Fisher Scientific). We used the same oligonucleotide primers and amplification protocols as in Borko et al., 2022, 2021. PCR product were purified and sent for bidirectional Sanger sequencing to Macrogen Europe laboratory (Amsterdam, Netherlands). Resulting chromatograms were edited, assembled and aligned in Geneious 11.0.3. (Biomatters Ltd).

**Table S1.** PCR primers used in the molecular analysis. All successfully amplified PCR products were sequenced by Macrogen Europe laboratory (Amsterdam, Netherlands), using the same amplification primers and bidirectional Sanger sequencing.

| **Marker** | **Primer** | **Primer sequence** | **Primer reference** | **Protocol reference** |
| --- | --- | --- | --- | --- |
| 28S rDNA, I | 28S lev2 | CAAGTACCGGTGAGGGAAAGTT | 9 | 6 |
|  | 28S des2 | GTTCACCATCTTTCGGGTC |  |  |
| 28S rDNA, II | 28S lev3 | GCCCTTAAAATGGATGGCGCT | 6 | 6 |
|  | 28S des5 | CCGCCGTTTACCCGCGCTT |  |  |
| 28S rDNA, III | 28S lev6 | TGTCAACAGTGATTGAACATGG | 12 | 12 |
|  | 28S des6 | GCGTTAAGCAGAAAAGAAAACTC |  |  |
| ArgK | ArgKin_F3 | CCCCTTCAACCCYTGYCTBACYGAGGC | 1 | 1 |
|  | ArgKin_R3 | GGVAGCTTRATRTGGACGGAGGC |  |  |
| Histone H3 | H3aF2 | ATGGCTCGGTACCAAGCAGAC | 5 | 6 |
|  | H3aR2 | ATRTCCTTGGGCATGATTGTTAC |  |  |
| EPRS | EPRS_1F | CAGGAAACAGCTATGACCGARAARGARAARTTYGC | 4 | 1 |
|  | EPRS_1R | TGTAAAACGACGGCCAGTTCCCARTGRTTRAAYTTCCA |  |  |
| HSP70 | HSP70F324 | GATCATCGCCAACGAYCAGGG | 7 | 7 |
|  | HSP70R960 | CGCTTGAAYTCYTGGATGAAGT |  |  |
| Cytochrome Oxidase I | LCO1490 | GGTCAACAAATCATAAAGATATTG | 2 | 3 |
|  | HCO2198 | TAAACTTCAGGGTGACCAAAAAAT |  |  |
| Cytochrome Oxidase II | LCO1490 | GGTCAACAAATCATAAAGATATTG | 10 | 10 |
|  | Spr1 | CGRTCTGTTARTAATATWGTAAT |  |  |
| 18S | 18sF | CCTAYCTGGTTGATCCTGCCAGT | 8 | 8 |
|  | 18sR | TAATGATCCTTCCGCAGGTT |  |  |
| ITS | F1 | TCCGAACTGGTGCACTTAGA | 11 | 11 |
|  | R1 | TCCAAGCTCCATTGGCTTAT |  |  |
|  | SF1 | CGCTGCCATTCTCACACTTA |  |  |
|  | SR1 | ACTCTGAGCGGTGGATCACT |  |  |
|  | SF2 | AAGGCTATAGCTGGCGATCA |  |  |
|  | SR2 | TCAGCGGGTAACCTCTCCTA |  |  |

11 Flot, J.-F., Wörheide, G., Dattagupta, S. Unsuspected diversity of *Niphargus* amphipods in the chemoautotrophic cave ecosystem of Frasassi, central Italy. BMC Evolutionary Biology 10, 171 (2010).

12 Ntakis, A., Anastasiadou, C., Zakšek, V., Fišer, C. Phylogeny and biogeography of three new species of *Niphargus* (Crustacea: Amphipoda) from Greece. Zoologischer Anzeiger - A Journal of Comparative Zoology 255, 32–46 (2015).

**Laboratory protocol by Eawag (Dübendorf, Switzerland)**

We used three pereopods for DNA isolation whenever possible. For small or poorly preserved specimens we used half or the whole body. To the dried body part we then added a 10% Chelex solution, including 10 μL proteinase K per sample (in total 160 μL Chelex-proteinase K solution per sample). The extraction reaction was incubated overnight at 55 °C. For the PCR reaction, we used the Multiplex PCR Kit from QIAGEN (Basel, Switzerland). We amplified the mitochondrial cytochrome oxidase I (COI) gene and two nuclear DNA gene fragments, namely parts of the 28S rRNA gene and parts of the internal transcribed spacer I and II (ITS). For COI, we used the primers LCO1490 and HCO2198 to amplify a 658 bp long fragment (Tab. S1). PCR reactions (25 μL) were obtained by mixing QIAGEN Multiplex PCR Mastermix (1x), Q Solution (0.5x), the two primers (0.5x each) and H2O, using 23 μL for each sample and 2 μL of the DNA extract. We used a PCR protocol with the following cycling settings: 95 °C for 15 min, 35 cycles at 94 °C for 30 s, 52 °C for 90 s and 72 °C for 60 s, and a final extension at 72 °C for 10 min before cooling to 10 °C. Considering 28S, we used the primer pair 28S-lev6 and 28S-des6 (Tab. S1), which amplified about 530 bp. For the PCR reactions (25 μL), we used the same quantities and concentrations as for COI, however, the DNA extract was diluted 1:5. The PCR reactions consisted of the following cycling settings: 95 °C for 15 min, 35 cycles at 94 °C for 30 s, 57 °C for 90 s and 72 °C for 60 s, and a final extension at 72 °C for 10 min before cooling to 10 °C. For ITS we amplified about 1,800 bp long fragments using the primers described by Flot et al. (2010) (Tab. S1) by mixing the same quantities and concentrations as for COI. An amount of 25 μL was used per sample and the PCR cycling settings were set to 95 °C for 15 min, 35 cycles at 94 °C for 30 s, 53 °C for 90 s and 72 °C for 180 s, and a final extension at 72 °C for 5 min before cooling to 10 °C. The PCR was conducted including the primers F1 and R1. We sent PCR products and corresponding primers to Microsynth AG (Balgach, Switzerland) for Sanger sequencing. For the sequencing reactions, the same primer pairs as for the PCR reactions were used. For ITS, internal primers SR1, SR2, SF1, and SF2 (Flot et al. 2010) were added, in addition to F1 and R1. Chromatograms were visually checked and edited in CodonCode Aligner 10.0.1 (CodonCode Corporation, Centerville, United States).

Colgan, D.J., McLauchlan, A., Wilson, G.D.F., Livingston, S.P., Edgecombe, G.D., Macaranas, J., Cassis, G., & Gray, M.R. (1998). Histone H3, U2 snRNA DNA sequences and arthropod molecular evolution. *Australian Journal of Zoology*, 46, 419–437.

Fišer, C., Zagmajster, M., & Zakšek, V. (2013). Coevolution of life history traits and morphology in female subterranean amphipods. *Oikos*, 122, 770–778.

Flot, J.-F., Wörheide, G., Dattaguptac, S. (2010). Unsuspected diversity of *Niphargus* amphipods in the chemoautotrophic cave ecosystem of Frasassi, central Italy*. BMC evolutionary biology* 10, p. 171.

Fujita, S.I., Senda, Y., Nakaguchi, S. & Hashimoto, T. (2001). Multiplex PCR using internal transcribed spacer 1 and 2 regions for rapid detection and identification of yeast strains. Journal of Clinical Microbiology, 39, 3617–3622

Folmer, O.M., Black, M., Hoeh, R., Lutz, R. & Vrijehoek, R. (1994). DNA primers for amplification of mitochondrial ytochrome *c* oxidase subunit I from diverse metazoan invertebrates. *Molecular Marine and Biotechnology*, 5, 304–313.

Simon, C., Franke, A. & Martin, A. (1991). The polymerase chain reaction: DNA extraction and amplification. Pp. 329–355 in: G.M. Hewitt, A.W.B. Johnson & J.P.W. Young (eds.), Molecular techniques in taxonomy. Springer-Verlag, Berlin.

Verovnik, R., Sket, B., & Trontelj, P. (2005). The colonization of Europe by the freshwater crustacean *Asellus aquaticus* (Crustacea: Isopoda) proceeded from ancient refugia and was directed by habitat connectivity. *Molecular Ecology*, 14, 4355–4369.

**Laboratory protocol by ULB (Brussels, Belgium)**

One or two pereopods of each specimen (sometimes more if specimens were juveniles) were used for DNA extraction, and the remaining parts of each specimen were stored in 96% EtOH at -20°C at ULB. Extraction of genomic DNA was performed using the NucleoSpin Tissue kit by MachereyNagel, following the manufacturer’s protocol. The eluted DNA was stored at 4°C until amplification, then long-term stored at -20°C. The following markers were PCR-amplified: (1) a fragment (between 986 and 998 bp long) of the nuclear 28S rRNA gene (Verovnik's fragment: external primers of Verovnik et al. 2005 complemented with internal primers of Flot et al. 2010): (ii) a 658 bp fragment (Folmer's fragment: Folmer et al. 1994, using the primers of Astrin & Stüben 2008) of the mitochondrial cytochrome c oxidase subunit I (COI); (3) the complete internal transcribed spacer (ITS) region (together with flanking portions of the 18S and 28S genes and including 5.8S, length between 1626 and 1815 bp) using the six primers described in Flot et al. (2010). PCR reactions for COI and 28S (10 μL) were obtained by mixing TaQ Green Mastermix (5 μL) the two primers (0.8 μL each at 10%), H2O (2.4 μL), and 1 μL of the DNA extract. PCR reactions for ITS (20 μL) were obtained by mixing TaQ Green Mastermix (10 μL) the two primers (1.6 μL each at 10%), H2O (4.8 μL), and 2 μL of the DNA extract. Direct sequencing was performed using the same primers as for amplification as well as internal primers (Flot et al. 2010); 28S, COI, and ITS PCR products were sent for bidirectional Sanger sequencing to Macrogen Europe (Amsterdam, The Netherlands).

| **Marker** | **Primer** | **Bases** | **Direction** | **Literature** | **Use** |
| --- | --- | --- | --- | --- | --- |
| COI | LCO1490-JJ | 5'-CHACWAAYCATAAAGATATYGG-3' | Forward | (Astrin & Stüben, 2008) | PCR + sequencing |
| COI | HCO2198-JJ | 5'-AWACTTCVGGRTGVCCAAARAATCA-3' | Reverse | (Astrin & Stüben, 2008) | PCR + sequencing |
| 28S | Niph15 | 5'-CAAGTACCGTGAGGGAAAGTT-3' | Forward | (Verovnik et al., 2005) | PCR |
| 28S | Niph16 | 5'-AGGGAAACTTCGGAGGGAACC-3' | Reverse | (Verovnik et al., 2005) | PCR |
| 28S | Niph15i | 5'-AGAGTCAAAAGACCGTGAAACC-3' | Forward | (Flot, unpublished | sequencing |
| 28S | Niph16i | 5’-GATTGGTCTTTCGCCCCTAT-3’ | Reverse | (Flot, unpublished) | sequencing |
| 28S | Niph20 | 5'-AAACACGGGCCAAGGAGTAT-3' | Forward | (Flot et al., 2010) | sequencing |
| 28S | Niph21 | 5'-TATACTCCTTGGCCCGTGTT-3' | Reverse | (Flot et al., 2010) | sequencing |
| ITS | Niph1 | 5′-TCCGAACTGGTGCACTTAGA-3′ | Forward | (Flot et al., 2010) | PCR + sequencing |
| ITS | Niph2 | 5′-TCCAAGCTCCATTGGCTTAT-3′ | Reverse | (Flot et al., 2010) | PCR + sequencing |
| ITS | Niph5 | 5′-ACTCTGAGCGGTGGATCACT-3′ | Forward | (Flot et al., 2010) | sequencing |
| ITS | Niph6 | 5′-CGCTGCCATTCTCACACTTA-3′ | Reverse | (Flot et al., 2010) | sequencing |
| ITS | Niph22 | 5′-AAGGCTATAGCTGGCGATCA-3′ | Forward | (Flot et al., 2010) | sequencing |
| ITS | Niph23 | 5′-TCAGCGGGTAACCTCTCCTA-3′ | Reverse | (Flot et al., 2010) | sequencing |

PCR conditions COI and 28S: 3min at 94°C, 30x (20sec at 94°C, 45sec at 50°C, 1min at 72°C), 2min at 72°C

PCR conditions ITS: 1min at 94°C, 40x (30sec at 94°C, 30sec at 53°C, 3 min at 72°C)

Astrin, J. J., Stüben, P. E. (2008). Phylogeny in cryptic weevils: molecules, morphology and new genera of western Palaearctic Cryptorhynchinae (Coleoptera: Curculionidae). Invertebrate Systematics, 22, 503–522.

Flot, J.-F., Wörheide, G., Dattagupta, S. (2010). Unsuspected diversity of *Niphargus* amphipods in the chemoautotrophic cave ecosystem of Frasassi, central Italy. BMC Evolutionary Biology, 10, 171.

Folmer, O., Black, M., Hoeh, W., Lutz, R., Vrijenhoek, R. (1994). DNA primers for amplification of mitochondrial cytochrome c oxidase subunit I from diverse metazoan invertebrates. Molecular Marine Biology and Biotechnology, 3, 294–299.

Verovnik, R., Sket, B., Trontelj, P. (2005). The colonization of Europe by the freshwater crustacean *Asellus aquaticus* (Crustacea: Isopoda) proceeded from ancient refugia and was directed by habitat connectivity. Molecular Ecology, 14, 4355-4369.
