## SupplementaryMaterial06 for "Evolutionary origin of morphologically cryptic species imprints co-occurrence and sympatry patterns"

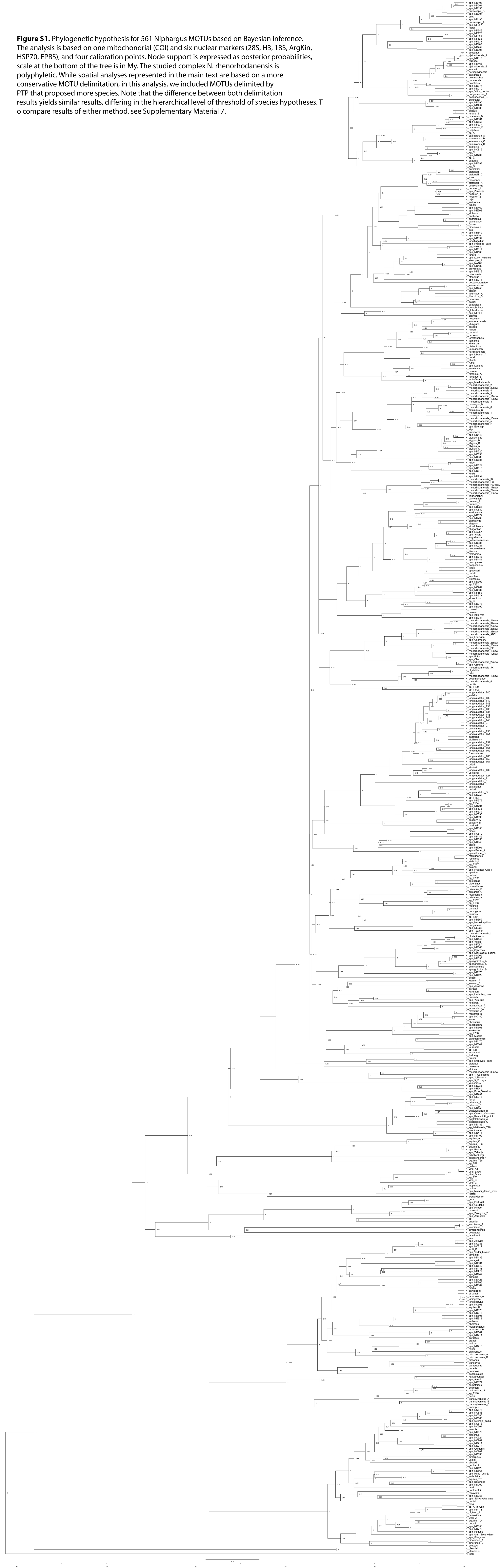

**Figure S1.** Phylogenetic hypothesis for 561 Niphargus MOTUs based on Bayesian inference. The analysis is based on one mitochondrial (COI) and six nuclear markers (28S, H3, 18S, ArgKin, HSP70, EPRS), and four calibration points. Node support is expressed as posterior probabilities, scale at the bottom of the tree is in My. The studied complex N. rhenorhodanensis is polyphyletic. While spatial analyses represented in the main text are based on a more conservative MOTU delimitation, in this analysis, we included MOTUs delimited by PTP that proposed more species. Note that the difference between both delimitation results yields similar results, differing in the hierarchical level of threshold of species hypotheses. T o compare results of either method, see Supplementary Material 7.
