## SupplementaryMaterial08 for "Evolutionary origin of morphologically cryptic species imprints co-occurrence and sympatry patterns"

*Running title: Sympatry and syntopy among cryptic species*

Teo Delić^1^, Špela Borko^1^, Ester Premate^1^, Behare Rexhepi^1^, Roman Alther^2,3^, Mara Knüsel^2,3^, Florian Malard^4^, Dieter Weber^5,6^, Fabio Stoch^7^, Jean-François Flot^7,8^, Cene Fišer^1*^, Florian Altermatt^2,3*^

^1^Department of Biology, Biotechnical Faculty, University of Ljubljana, Jamnikarjeva 101, Ljubljana, Slovenia

^2^Department of Aquatic Ecology, Eawag, Swiss Federal Institute of Aquatic Science and Technology, Überlandstrasse 133, 8600 Dübendorf, Switzerland

^3^Department of Evolutionary Biology and Environmental Studies, University of Zurich, Winterthurerstrasse 190, 8057 Zurich, Switzerland

^4^Univ Lyon, Université Claude Bernard Lyon 1, CNRS, ENTPE, UMR 5023 LEHNA, F-69622, Villeurbanne, France

^5^Senckenberg Deutsches Entomologisches Institut, Eberswalder Straße 90 15374 Müncheberg, Germany

^6^Musée National d'Histoire Naturelle de Luxembourg, 25 Rue Münster, L-2160 Luxembourg, Luxembourg

^7^Evolutionary Biology & Ecology, Université libre de Bruxelles (ULB), Brussels, Belgium

^8^Interuniversity Institute of Bioinformatics in Brussels – (IB)2 , Brussels, Belgium

**Supplementary Figures S2 – S7**

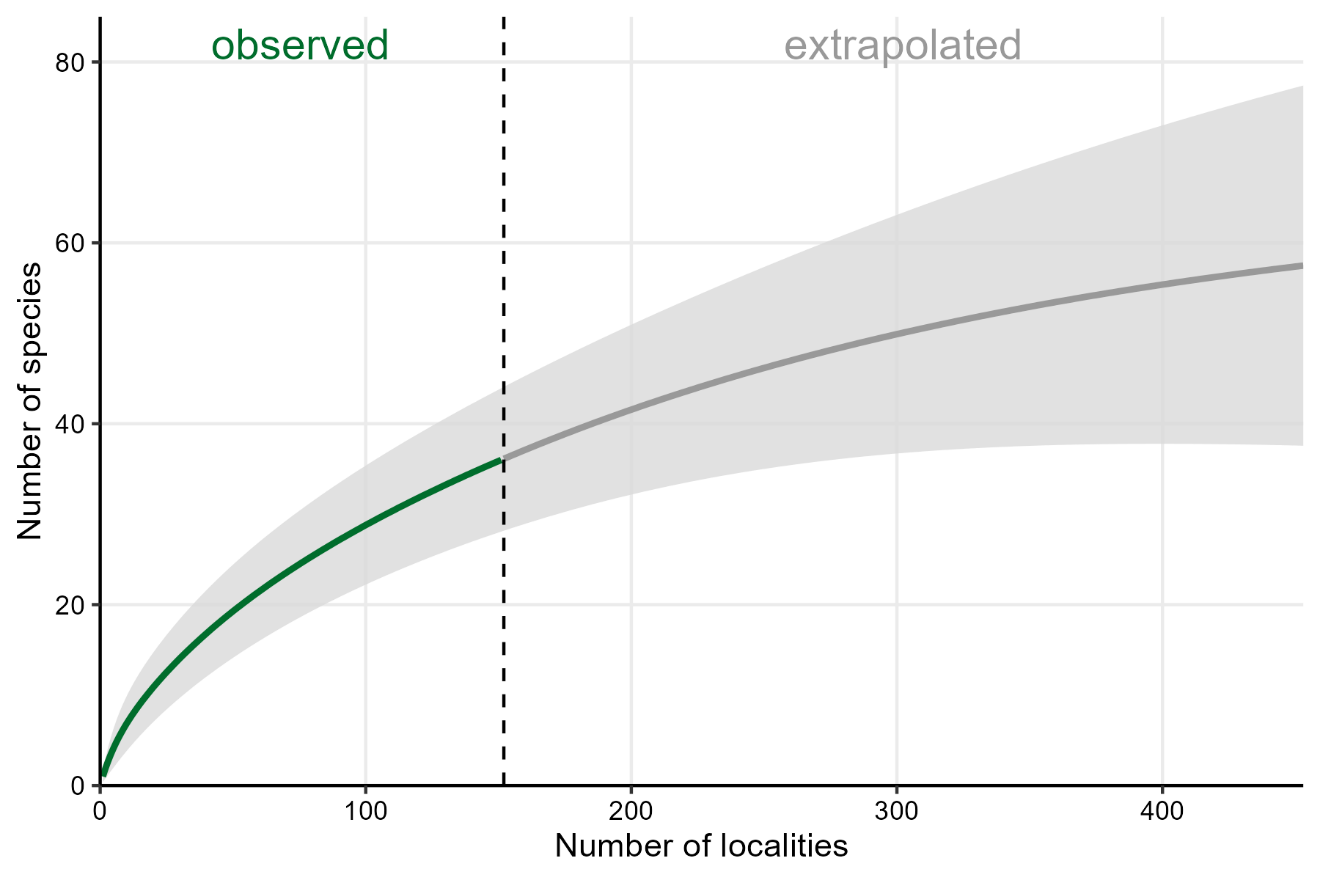

**Figure S2.** Rarefaction curve. We analyzed the completness of our taxon sampling usig EstimateS and MOTUs of the studied species complex *N. rhenorhodanesis*, delimited by a more conservative method ASAP. Sampling unit was a locality (151 localities, dashed vertical line). The analysis is extrapolated to threefold sampling effort and is weakly leveling off. Shaded area represent 95 % confidence interval.

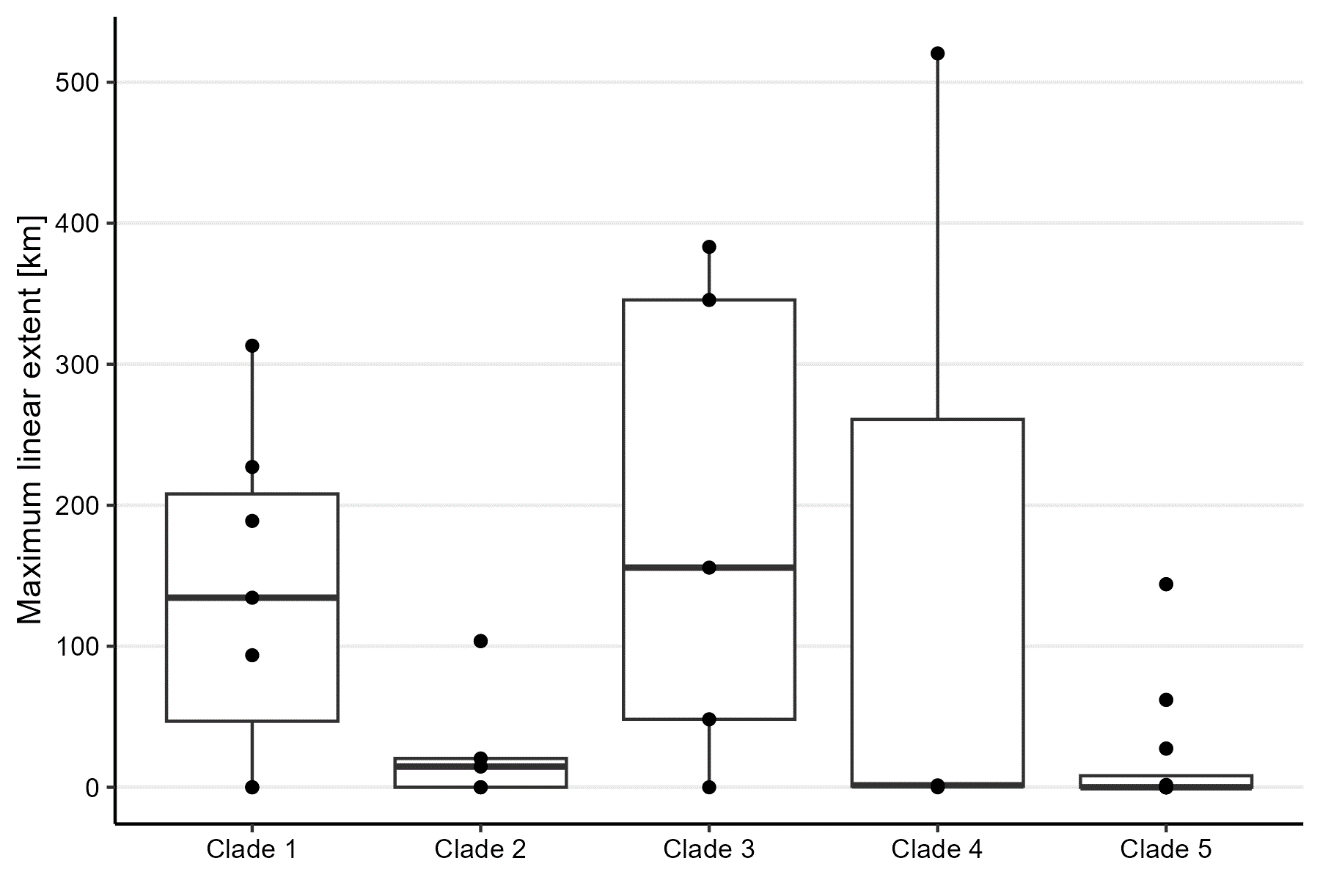

Figure S3. Range sizes of the MOTUs from five main clades of *N. rhenorhodanensis* species complex. Range sizes are expressed as maximum linear extent of the longest diagonal of the range. Each clade comprises single-site species and large range species.

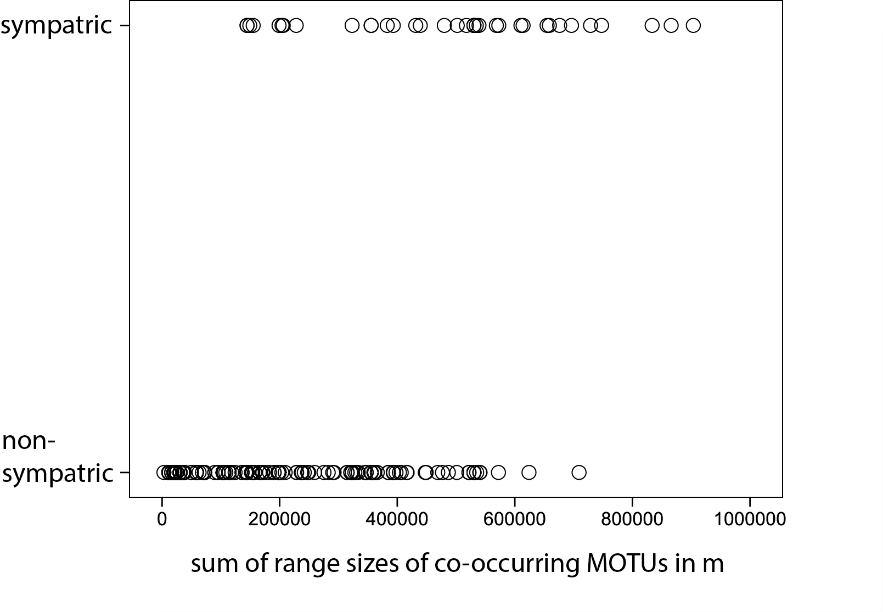

**Figure S4.** Distribution of sympatric and non-sympatric MOTU pairs with respect to the range size. The X axis shows a sum of range sizes estimated as maximum linear extent, and were used as predictors in generalized linear model. Response variable was a binary, sympatric versus nonsympatric, and the mode was statistically significant (see main text).

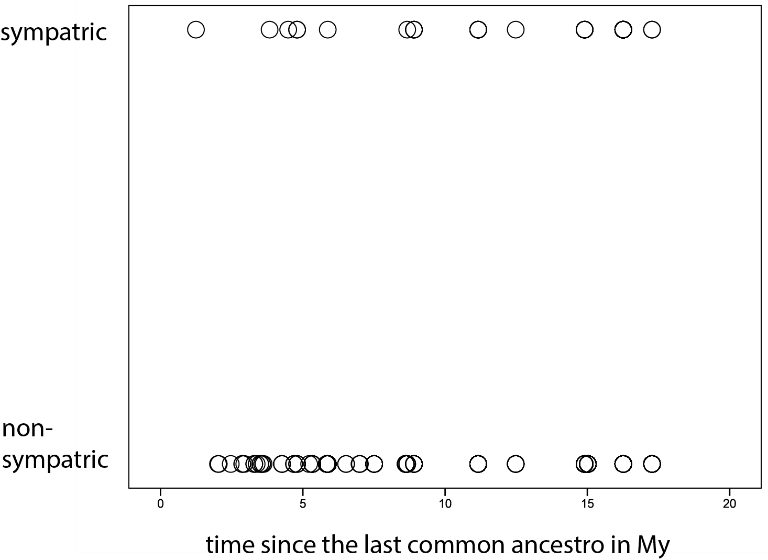

**Figure S5.** Distribution of sympatric and non-sympatric MOTU pairs with respect to the time since the last ancestor, estimated from the phylogeny. Using these data, we ran generalized linear model with age of a pair as predictor, and sympatry versus non-sympatry as a binary response variable. The model was non-significant (see main text).

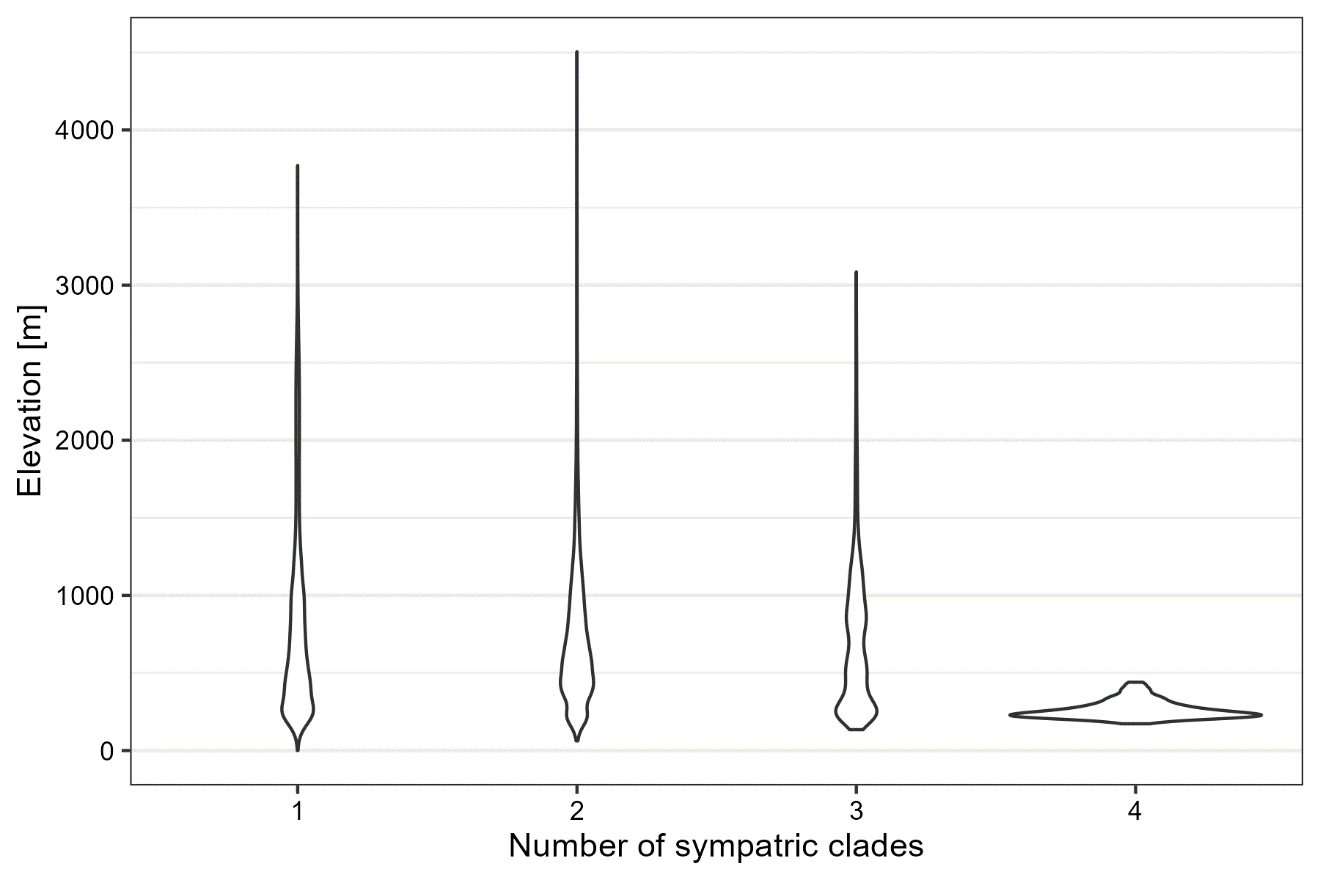

**Figure S6**. Number of sympatric clades, with respect to elevational range. We quantified the altitudes at resolution 1 x 1 km of regions, where in sympatry live two, three or four clades. Here, we present full altitudinal range, whereas in the main text we truncated plots at the elevation of 1500 m a.s.l.

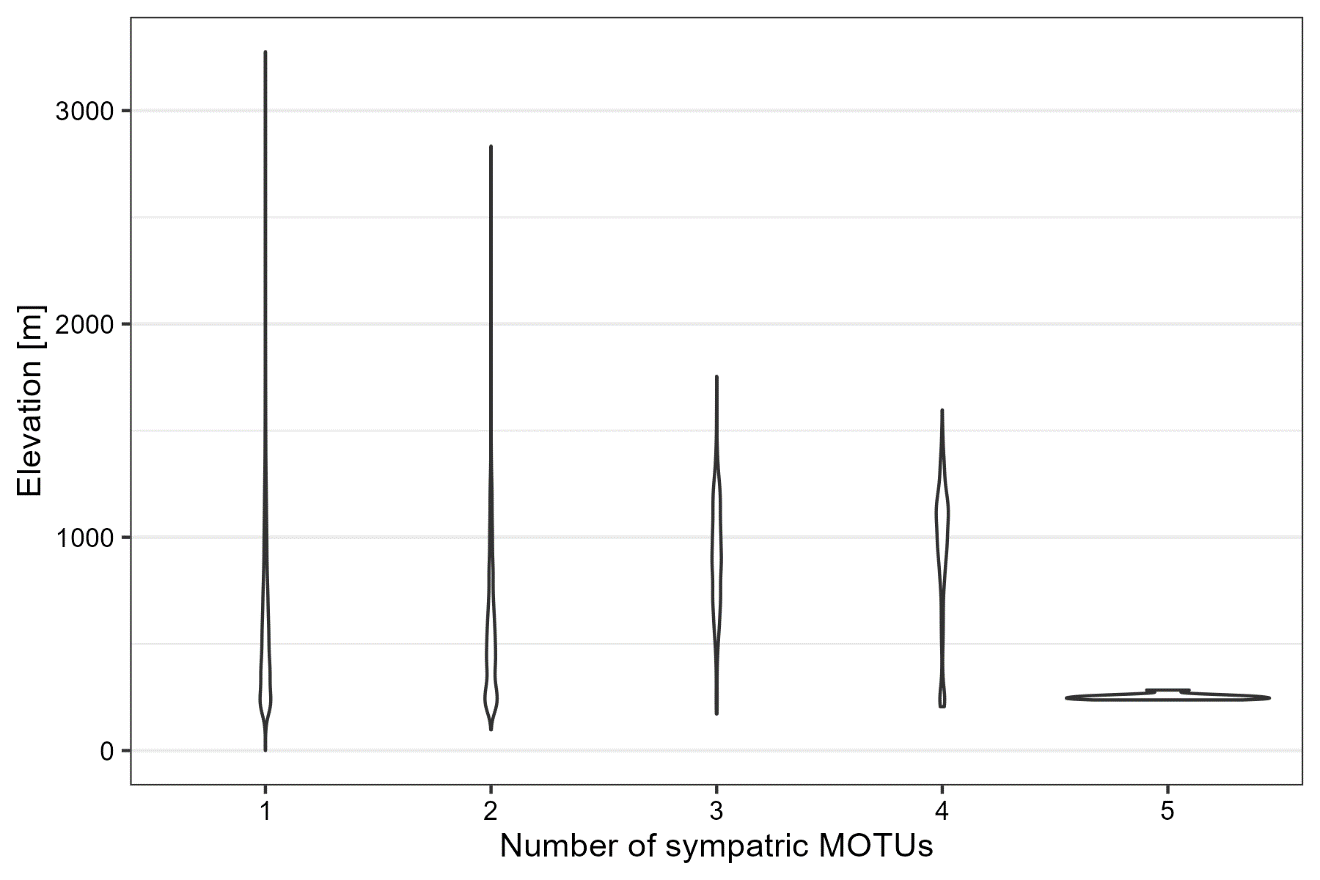

Figure S7. Number of sympatric MOTUs, with respect to elevational range. We quantified the elevations at resolution 1 x 1 km of regions, where in sympatry live two, three or four clades. Here, we present full elevational range, whereas in the main text we truncated plots at the elevation of 1500 m a.s.l.
